# Pitfalls in the Assessment of Brain-Machine Interfaces Using Information Transfer Rate

**DOI:** 10.1101/205013

**Authors:** Mikhail A. Lebedev, Po-He Tseng, Peter J. Ifft, Dennis Ochei, Miguel A.L. Nicolelis

**Author notes:** **Corresponding Authors:** Mikhail A. Lebedev, PhD,; Miguel A. L. Nicolelis, MD, PhD.

## Abstract

Information transfer rate (ITR), measured in bits/s, can be applied to evaluate motor performance, including the capacity of brain-machine interfaces (BMIs) to control external actuators. In a 2013 article entitled “Transfer of information by BMI” and published in *Neuroscience*, Tehovnik and his colleagues utilized ITR to assess the performance of several BMIs reported in the literature. We examined these analyses closely and found several fundamental flaws in their evaluation of ITR. Here we discuss the pitfalls in Tehovnik’s measurements of ITR, as well as several other issues raised in “Transfer of information by BMI”, including the claim that BMIs cannot be a reasonable option for paralyzed patients.

**Highlights:**

- Information transfer rate is discussed for BMI experiments, where subjects reach to targets.
- Task settings, not just the number of possible targets, are important to calculate information correctly.
- Active tactile exploration can be quantified as information transfer, but the number of targets is insufficient for such quantification.
- Information transfer rate increases with the number of neural recording channels.
- For practical applications, improvement in quality of life is essential, not information transfer rate per se.

## Introduction

In a 2013 review paper entitled “Transfer of information by BMI” (Tehovnik, Woods et al. 2013), Tehovnik and his colleagues claimed to have performed a systematic evaluation of ITR for both invasive and noninvasive BMIs reported in the literature. They restricted their analysis to center-out experiments because, in their opinion, these datasets could be analyzed using a straightforward, uniform approach. Although their analyses might appear to be credible at first sight, we found them flawed, mostly because the cited BMI results were interpreted incorrectly.

## Information Metric for Center-Out Task

Tehovnik and his colleagues calculated ITR for a center-out task using a metric previously used by Wolpaw et al. (Wolpaw, Birbaumer et al. 2000, Wolpaw, Birbaumer et al. 2002). Wolpaw et al.’s approach is not original. An earlier paper by Sakitt (Sakitt 1980) developed a method for calculating ITR for an experiment where a subject performs reaching movements to screen targets (Fig. 1). Under these conditions, information contained in the targets, *H(u)*, is reproduced, with some information loss, by motor responses that carry information *H(v)*. If *N*_*i*_ is the number of times stimulus *u*_*i*_ is presented, *N*_*j*_ is the number of times the response *V*_*j*_ is produced, and *N*_*ij*_ is their joint count (also called stimulus-response matrix) then

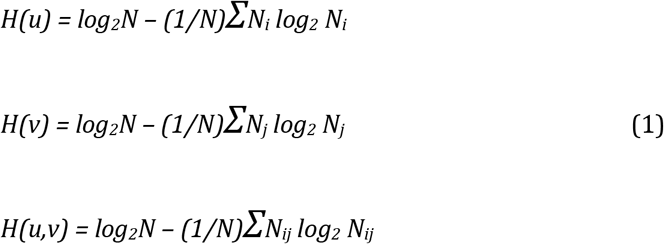

where *N* is the total number of trials and *H(u,v)* is joint information. Mutual information (or information transmitted by the channel), *T(u,v)*, is given by Shannon’s equation (Shannon and Weaver 1964):

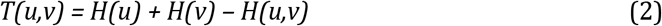

**Figure 1.**
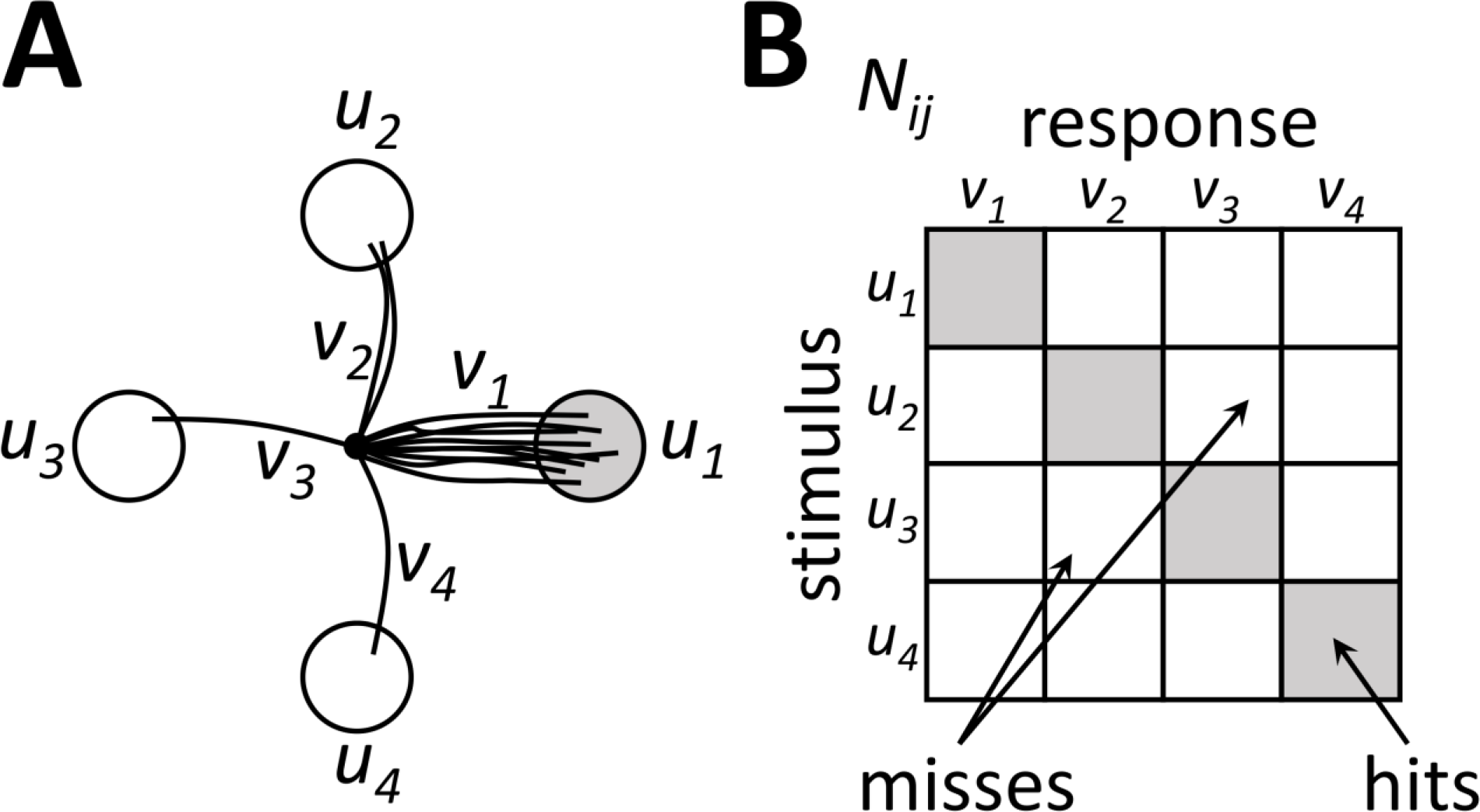
Schematics for information transfer model proposed by Sakitt (1980). (A) An illustration of a center-out task with 4 target locations (*u*_*1-4*_). On the behavioral trials when stimulus *u*_*1*_ is presented, many responses are correct (*v*_*1*_) but a few errors are made (*v*_*2*_, *v*_*3*_, and *v*_*4*_). Errors decrease mutual information, *T(u,v)*, the quantity that can be used to evaluate the task performance. (B) Stimulus-response matrix, *N*_*ij*_, represents trial counts for different stimulus-response combinations. The matrix diagonal represents correct responses (hits), whereas the off-diagonal elements represent incorrect responses (misses).

Sakitt’s algorithm was employed by Georgopoulos and Massey (Georgopoulos and Massey 1988) to estimate the ITR for neuronal samples collected in motor cortex and cortical area 5. In these experiments, monkeys were engaged in an arm reaching task. Georgopoulos and Massey utilized a population vector method to extract directional information from the neuronal activity. Additionally, they calculated the ITR for center-out reaching movements performed by human subjects. The estimates for monkey neurons and human subjects were comparable (around 5 bits per trial or 1 bit/s).

Wolpaw et al. (Wolpaw, Birbaumer et al. 2000, Wolpaw, Birbaumer et al. 2002) used a simplified derivation of *T(u,v)*, which they obtained from Pierce (Pierce 1980). This derivation assumes equal probabilities of correct responses for different stimuli. *T(u,v)* is calculated as:

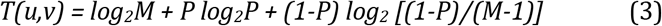

where *P* is the proportion of correctly performed trials, and *M* is the number of choices (e.g. the number of targets on the screen from which a subject selects one).

## Pitfalls in Tehovnik’s Measurement of Information Transfer

Although equation 3 is, in principle, applicable to ITR in BMIs, the way in which Tehovnik and his colleagues interpreted this method was flawed. A relatively minor error was to assume that in all BMI experiments the probability of correct response is the same for different target locations. However, the major flaw was related to the assumption that the knowledge of the number of screen locations, *M*, where targets *could* appear was sufficient to calculate the ITR. According to Tehovnik and his colleagues:

> *“To determine the bits per trial of a particular subject (whether human or monkey) in a BMI study, one needs two pieces of information: the percent-correct score and the number of targets.”*

In reality, these authors’ selection of these “two pieces of information” resulted in erroneous results. Essentially, they missed the point that the number of targets (*M* in equation 3) should represent real choices on each trial. Such choices could be achieved, for example, by showing *M* targets on the screen, instructing the subject to reach toward one of them, and cancelling the trial if the subject reaches to an incorrect target. Even if all potential targets were not shown on the screen, responses could be deemed erroneous when the subject intersects the location of an incorrect target. The experiments cited by Tehovnik and his colleagues did not have such a provision. In these experiments, only one target was shown on the screen on each trial, and subjects were allowed to cross any screen location until they reached that target. Such a single target condition is like a telephone dial pad with just one button.

Among the cited literature, Tehovnik and his colleagues referred to the study of Ganguly and Carmena (Ganguly and Carmena 2009) but erroneously asserted that monkeys selected from eight targets in these experiments. In this study, a lone peripheral target was present on the screen on each trial (Fig. 2A). (This is perhaps not explained clearly in the paper, but is illustrated in the 28^th^ minute of their video: https://youtu.be/29wFC3Wgbgc.) The fact that target location on each trial was chosen from eight preset locations was irrelevant because monkeys were not penalized for crossing wrong target locations with the cursor. (Such crossings can be seen in Fig. 2D of Ganguly and Carmena, 2009.) Trials were only canceled if a monkey failed to acquire the target within a 10-s time limit. Accordingly, there were only two possible outcomes in this behavior: target acquisition or a timeout, not eight as Tehovnik assumed. Note, however, that the possibility of two outcomes was not equivalent to 1 bit of information because a failure to acquire the target did not represent a meaningful selection. (It would if, for example, the monkey task was to report the presence or absence of the target.) A straightforward application of Sakitt’s model to these conditions would return an information content equal to zero. Indeed, for a single target present on the screen and no penalty for crossing incorrect target locations, rewarded performance could have been achieved even with completely random movements of the cursor, provided that the random walk occurred fast enough to cover the entire screen before the timeout occurred, but also not too fast because the cursor had to stay over the target for a required minimal time of 200–500 ms. This simple consideration invalidates Tehovnik’s calculations because random walk cannot transmit any information.

**Figure 2.**
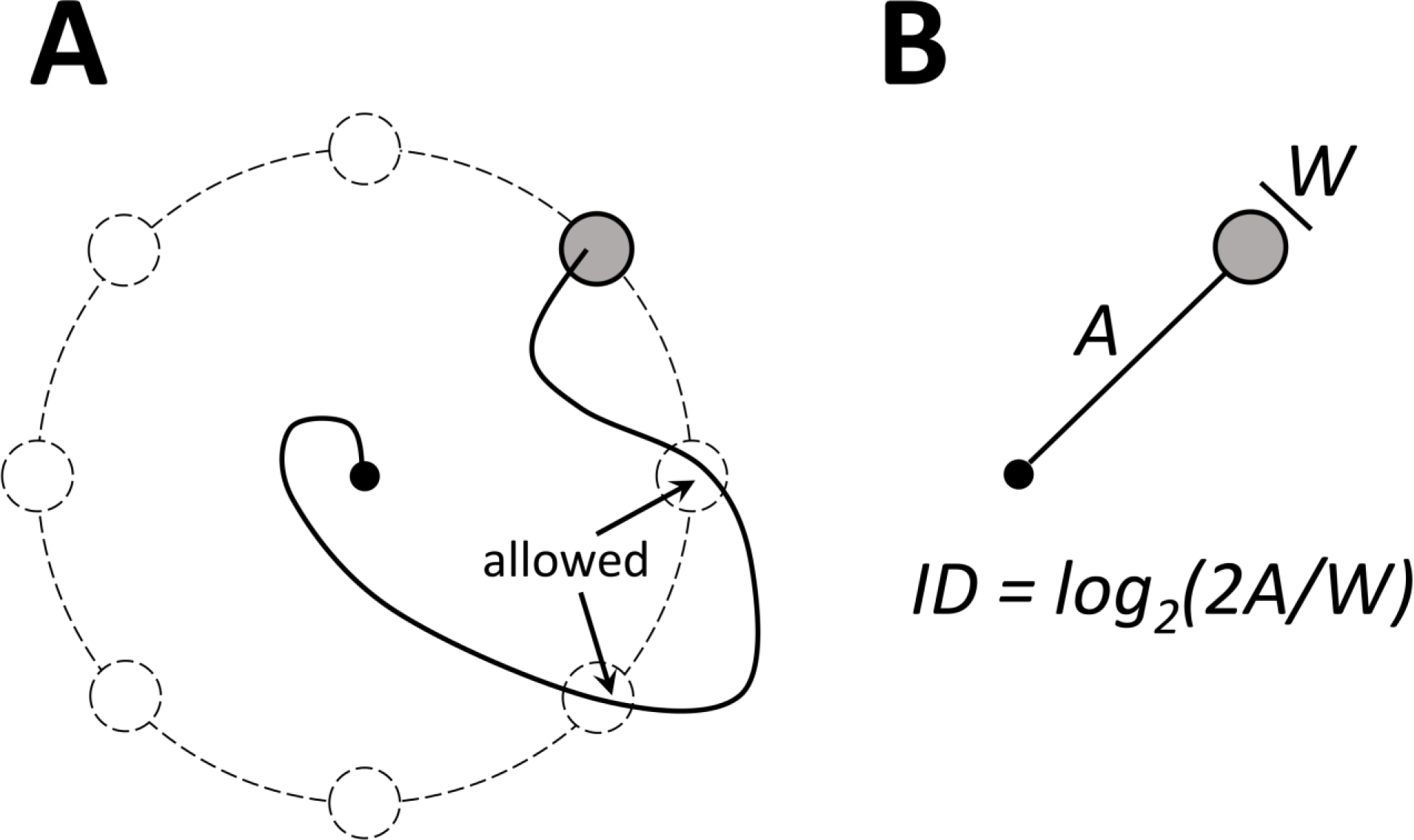
A common BMI paradigm with a single target shown on the screen. (A) A typical center-out task. A single trial is illustrated, where the target of movement (highlighted in gray) was chosen from eight possible locations (shown in dashed lines). Only one target is shown on the screen on each target. Although for the illustrated trial the cursor trajectory crossed two potential target locations, the trial was considered successful because the cursor eventually entered the visible target. (B) Fitts’ index of difficulty, *ID*, measured in bits and calculated based on the distance to target, *A*, and target width, *W*. This metric can be used as an alternative to Sakitt’s model to assess ITR when there is only one target shown on the screen and crossing other potential target locations is allowed.

To further illustrate the error in Tehovnik’s interpretation of parameter *M* in equation 3, they overlooked the fact that in some of the very studies that they quoted (e.g. O’Doherty et al. (O’Doherty, Lebedev et al. 2011)), targets were selected from an infinite number of potential locations, since the target angle was a randomly generated continuous variable. For an infinite number of targets, ITR approaches infinity – an obviously incorrect assessment.

Similarly, Tehovnik incorrectly presumed that Gilja and his colleagues (Gilja, Nuyujukian et al. 2012) had their monkeys select from eight available targets. Here again, there was only one target on the screen on each trial (see their Supplementary videos 1 and 2). Because of this task design, Gilja et al. could not apply Sakitt’s model and instead used Fitts’ metric (Fitts and Deininger 1954, Fitts and Peterson 1964) to compute the information:

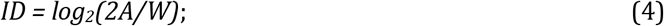

where *ID* is index of difficulty, measured in bits, *A* is distance to target, and *W* is target width (Fig. 2B). This formulation of information per trial, however, is not without problems by itself. It is acceptable for behaviors where subjects clearly perform reaching movements, but *ID* would be also non-zero for a random walk.

In addition to incorrectly interpreting monkey BMI experiments, Tehovnik misunderstood the behavioral task of Wolpaw and MacFarland (Wolpaw and McFarland 2004), the study conducted in human subjects. In these experiments, there was one target per trial, and an incorrect response occurred only in the case of a timeout, not when a subject intersected a wrong target location (see their Supporting Movie 1). The authors used coefficient of determination, *R*^2^, instead of ITR to quantify their results.

Furthermore, while equations 3 and 4, if applied correctly, are suitable for single decision classification tasks, neither of them are well suited to characterize continuous movements generated using a BMI. Indeed, trajectories carry more information than an average number of correctly completed trials and their average duration. ITR for continuous trajectories could be assessed, for example, by compute the entropy over all trajectories, and then subtracting the conditional entropy for the trajectories belonging to given targets. Such a calculation can be done by binning cursor positions using a grid and measuring the time-dependent occupancy probability for each bin, for all targets and for given targets.

Finally, equations 3 and 4 do not take into account fundamental requirements of BMI tasks, like accurately acquiring a target under a time limit. Of particular importance is the time required to hold the central and peripheral targets – one of the major contributors to task difficulty. Hence, even if a discrete method were applied correctly, the real ITR would have been estimated only approximately. Additionally, changing target hold duration and other task intervals would result in changes to the ITR calculated as information per trial divided by trial duration.

## Applying Sakitt’s Approach to Single-Target Reaches

Although Tehovnik and his colleagues measured ITR incorrectly, Sakitt’s model could be adapted to reaching movements towards a single target, when no alternative choices are shown on the screen and there is no penalty for crossing an incorrect target location. This can be done, for example, by generating the matrix *N*_*ij*_ by counting an intersection of a wrong target location as a behavioral choice. Additionally, mutual information represented by the stimulus and motor response could be estimated continuously, using a classifier that derives choices from where the cursor is moving (Fig. 3). In agreement with Sakitt’s model, such a classifier considers *M* possible targets, and deems one of them the chosen target. The chosen target could be derived from the cursor kinematics in different ways. One simple way to conduct such decoding is to find the target with the minimal distance to the cursor (Fig. 3). A more sophisticated method could incorporate cursor velocity. The basic idea here is that the BMI output carries information even before the final goal is accomplished. Once the cursor starts to move toward the target, target location can be predicted from the cursor trajectory. Predicting target location in advance is a well-known mathematical and engineering problem. Examples include calculation of projectile motion from the initial velocity (Newton 1687), estimation of the mobile user’s next mobile base station from the user’s location, heading, and altitude (Pathirana, Savkin et al. 2004), and recovery of video tracking of moving objects after an occlusion (Rosales and Sclaroff 1999). We suggest that a similar approach could be applied to analyze time dependence of information transfer by BMIs. With this method, transmitted information can be represented as function of time.

**Figure 3.**
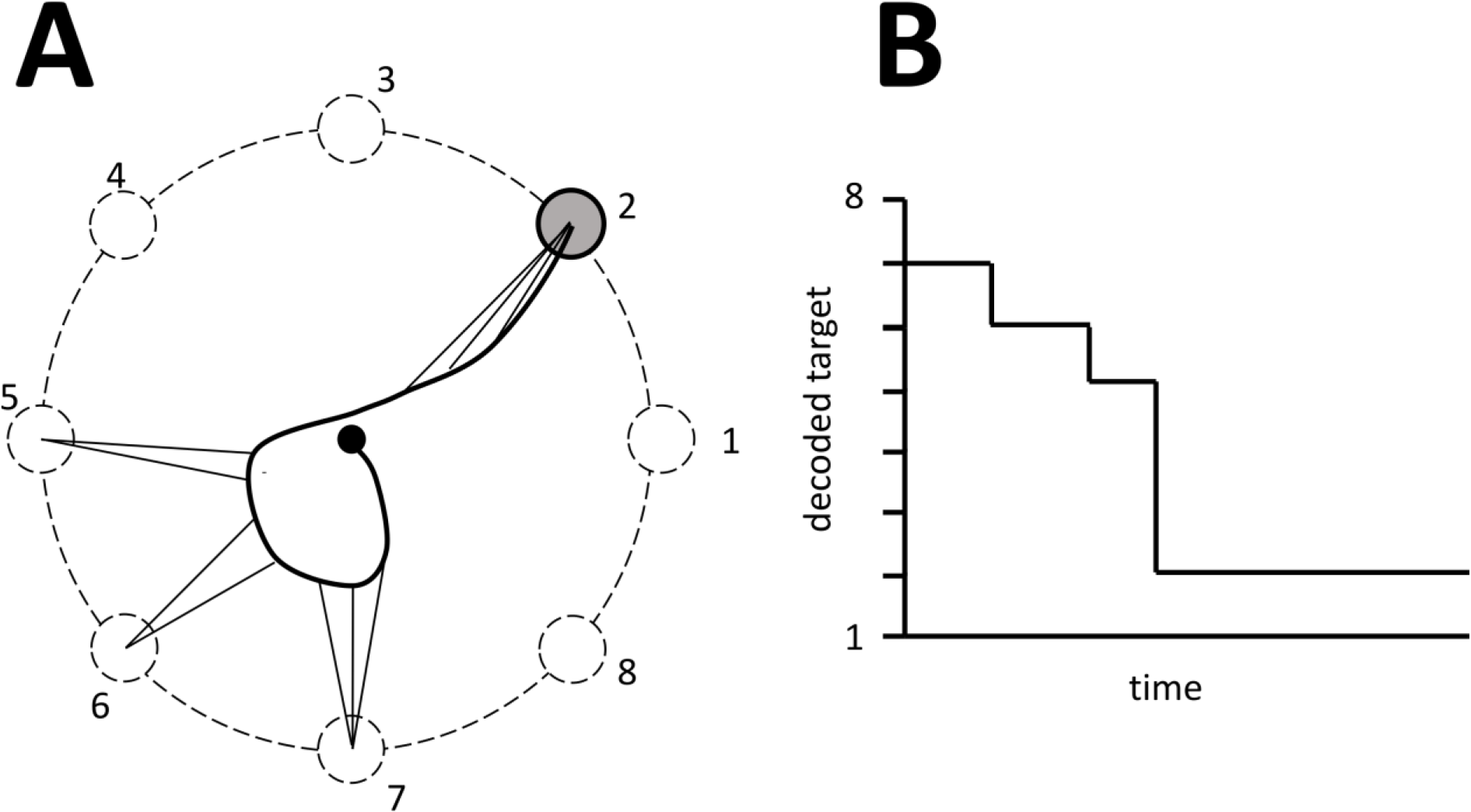
Continuous decoding of target location from cursor position. (A) Schematics of the decoding. As the cursor travels from the center to the screen target (highlighted in gray), intended target location is continuously decoded from the cursor position as the potential target with the minimal distance to the cursor (shown by lines connecting different points of the trajectory to the decoded targets). (B) Decoded target as a function of time.

## “Five well isolated cells” versus Multielectrode Methods

In addition to the attempt to assess ITR in previous BMI studies, Tehovnik and his colleagues expressed several critiques regarding the state of BMI field. Among these commentaries, they claimed that neurophysiologists have achieved a much higher ITR with traditional single-unit recordings, where just a few neurons were sampled at a time, compared to the recent BMI studies where neurons were sampled in large numbers. In particular, they claimed that ITR was 0.67 bit/s in Schmidt et al. (Schmidt, McIntosh et al. 1978), and only 0.03-0.17 bit/s in O’Doherty (O’Doherty, Lebedev et al. 2011).

From the methods description of the first cited study (Schmidt, McIntosh et al. 1978), it is not clear whether or not moneys were allowed to intersect incorrect target locations, so we are unable to verify the ITR estimation. As to the study of O’Doherty and his colleagues (O’Doherty, Lebedev et al. 2011), Tehovnik and his colleagues produced a totally erroneous characterization of ITR. The paper reported a brain-machine-brain interface (BMBI) that enabled bidirectional information transfer between the brain and an avatar hand (Fig. 4). Although three targets were present on the screen in each trial, this number had nothing to do with *M* in equation 3, simply because monkeys were not given any initial instruction of the correct target. Therefore, this three-target task did not match Sakitt’s model at all. The behavioral task was an active exploration of targets guided by an artificial tactile sensation. Monkeys consecutively reached to each of the three targets with an avatar hand. When the avatar hand was put over a target, electrical microstimulation was applied to the primary somatosensory cortex, mimicking an artificial texture. The monkeys’ task was to find the target with a particular artificial texture and to hold the avatar hand over that target for at least 500 ms to obtain a reward. Acquisition of an incorrect target resulted in trial cancellation. Thus, the task was far more complex than pointing to a highlighted target when several others are present on the screen (Sakitt 1980). In principle, information-theoretic approach could be applied to the data of O’Doherty and his colleagues. For example, the processing of artificial sensory feedback could be modeled as a 1-bit choice: continue holding the target or leave. The ITR would then depend on the time spent making this decision. Given that in the study of O’Doherty average target exploration time was approximately 500 ms, ITR for this type of information transfer was close to 2 bits/s and not 0.03-0.17 bit/s as claimed by Tehovnik. A more complex evaluation of BMBI information transfer should take into account how information accumulates during active-exploration trials and how this information affects the subject’s decision regarding the sequence in which different targets are being exploration. For example, O’Doherty et al. found that monkeys rarely revisited unrewarded artificial textures, but often reexamined the rewarded ones.

**Figure 4.**
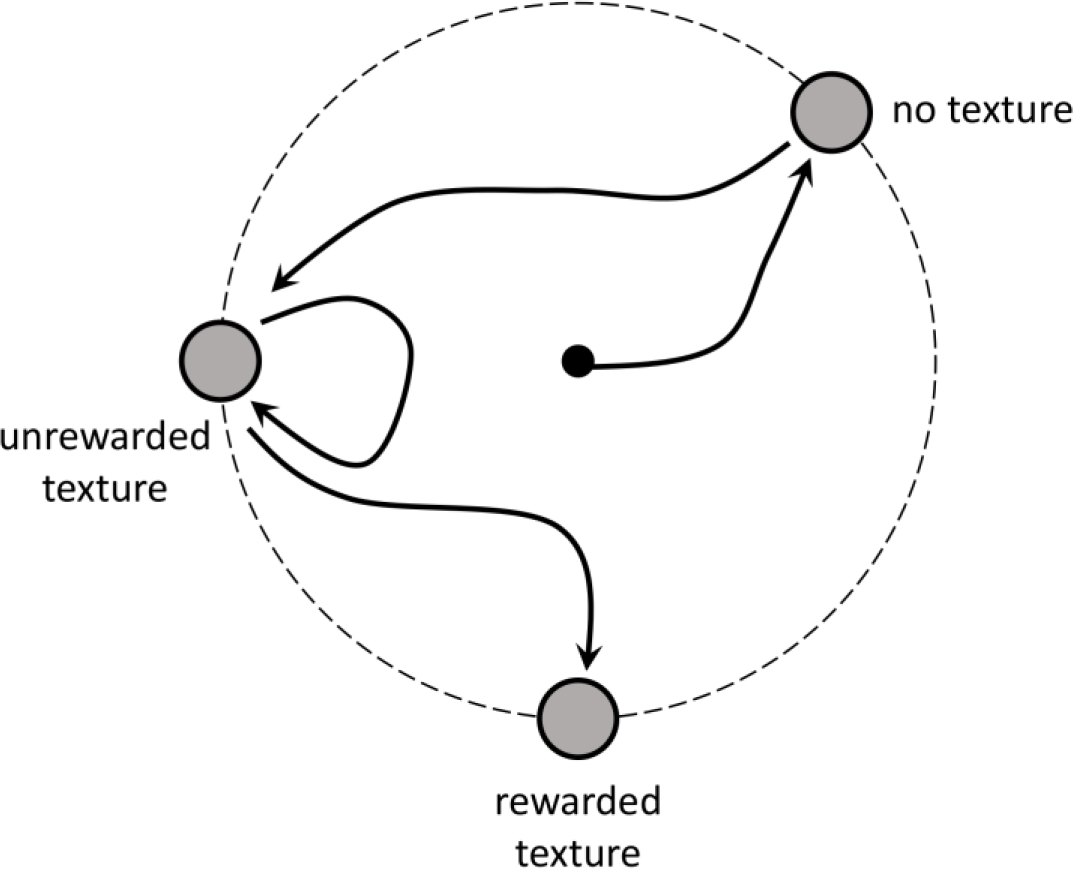
Active tactile exploration using a BMBI (O’Doherty et al. 2011). In this experiment, BMI output (position of an avatar hand on the screen) was derived from neuronal ensemble activity recorded in monkey motor cortex, while artificial tactile feedback was provided by microstimulation of somatosensory cortex. Although three targets were shown on the screen, Sakitt’s model is unapplicable because the behavioral task was different from the one shown in Fig.1. All targets were visually identical, and there was no advanced information regarding the correct target. To find the correct target, monkeys performed an active exploration: they placed the avatar hand over the targets to feel the artificial texture indicated by microstimulation. The animals discriminated three virtual textures: (1) no microstimulation, (2) unrewarded texture represented by 5-Hz packets of microstimulation, and (3) rewarded texture represented by 10-Hz packets. The figure illustrates a single trial, where the monkey first explored the no-texture target. Next, the monkey moved to the unrewarded-texture target and explored it twice by placing the avatar hand over that target, moving it out, and placing the avatar hand over the target again. Finally, after establishing that the first two targets were incorrect ones, the monkey moved the avatar hand to the third target, felt the rewarded artificial texture, and continued to hold the target to obtain a reward. Clearly, information transfer was quite complex in this experiment: monkeys obtain information by actively exploring the targets, and even hitting a wrong target contributed to this information accumulation and to achieving the eventual goal.

To explain why ITR was higher in Schmidt’s study than in O’Doherty’s study, Tehovnik claimed that “5 well isolated cells using movable electrodes can generate as much information as 100 poorly isolated cells using fixed electrodes”. However, in Schmidt’s study (Schmidt, McIntosh et al. 1978), the electrodes were actually implanted and fixed. Additionally, movable electrodes introduce several problems: (i) searching for task-related neurons by moving an electrode back and forth is harmful to the brain tissue, (ii) the method is successful only occasionally and often results in no single-units being isolated, and (iii) the method allows holding a unit for approximately half an hour, after which the unit is usually lost. Therefore, a fishing expedition for “star cells” is hardly an approach that would have any practical clinical significance for BMIs. Even Mulliken and his colleagues (Mulliken, Musallam et al. 2008), whose study Tehovnik et al. cited as an example of success, reported that they were capable of holding units for only 1-2 hours with their movable array. In a recent paper, our laboratory has demonstrated that high quality neuronal recordings can last over 5 years in monkeys implanted with our multi-electrode arrays and cubes (Schwarz, Lebedev et al. 2014).

Tehovnik and his colleagues also claimed that the ITR begins to saturate when the number of neurons increases beyond 50. This misconception is related to a flawed procedure where, after the subpopulation sample size approaches the total number of neurons in the population, there is not enough statistical variability in the data to draw any conclusion on whether the ITR is growing or has saturated (Lebedev 2014). Additionally, a closer inspection of the articles that Tehovnik and colleagues used to support their claim does not reveal any clear example where a neuron dropping curve would saturate at a fixed level. Rather, each of these articles shows a continuous improvement in decoding accuracy with the population size, and one quoted study specifically suggests that adding neurons results in a steady improvement of BMI accuracy (Wessberg, Stambaugh et al. 2000). We have previously explained that the dependency of decoding accuracy on the number of recorded neurons can be described by a logarithmic function (Lebedev and Nicolelis 2011, Ifft, Shokur et al. 2013), and a slower growth of the curve for large neuronal populations is consistent with this model.

It should be also noted that large populations of brain neurons allow simultaneous decoding of multiple behavioral parameters (Fitzsimmons, Lebedev et al. 2009) in situations where complex BMI controls are needed, such as bipedal walking (Fitzsimmons, Lebedev et al. 2009), reaching and grasping (Carmena, Lebedev et al. 2003, Hochberg, Bacher et al. 2012, Collinger, Wodlinger et al. 2013, Ajiboye, Willett et al. 2017), and bimanual operations (Ifft, Shokur et al. 2013). We have already discussed these issues in detail elsewhere (Nicolelis and Lebedev 2009) and urge readers interested in a discussion about the potential of BMI performance to read these original research papers as well as a recent review (Lebedev and Nicolelis 2017).

## Single Cells versus EEG

Tehovnik and his colleagues attempted to compare the performance of BMIs based on single cell recordings with those based on EEG. They cited the work by Wolpaw and McFarland (Wolpaw and McFarland 2004) in which four human subjects performed a center-out task using a continuous control of cursor movements by EEG signals. As mentioned above, Tehovnik assumed that subjects acquired one target from an array of eight, but in actuality there was a single target per trial, and crossing the other potential target locations was allowed and often happened. Building on these erroneous arguments, Tehovnik confronted Wolpaw and McFarland’s original claim that the performance of their BMI was comparable with BMIs based on single-cell recordings. According to Tehovnik, single-cell BMIs perform better – the conclusion that actually may be accidentally correct; but a correct analysis of ITR is needed, of course.

## Plasticity and BMI

Tehovnik and his colleagues suggested that an occurrence of unidirectional tuning of cortical cells cannot be good for the generation of a BMI’s optimal output. The justification of this statement is unclear. As follows from simple geometrical considerations, just two tuning directions (e.g. X and Y axes or any other basis) would be sufficient for a two-dimensional cursor control. Furthermore, Tehovnik claimed that increased synchronization between cells during BMI control (Carmena, Lebedev et al. 2003) would damage BMI performance. Recently, we showed that synchronization of neurons during BMI control is a transient effect that characterizes early BMI sessions (Ifft, Shokur et al. 2013). As monkeys improve in BMI control, synchronous firing subsides. While an analysis of all these details is beyond the scope of this article, we emphasize that any claim about BMI accuracy should involve a quantitative analysis.

Tehovnik also argued that it would be hard to build a stable BMI because the directional tuning of brain neurons fluctuates in time. However, Carmena and his colleagues (Carmena, Lebedev et al. 2005), Rokni et al. (Rokni, Richardson et al. 2007) and Huber et al. (Huber, Gutnisky et al. 2012) have demonstrated that representation at the population-level can be stable even when the tuning is unstable at the individual neuron level. Carmena (Carmena, Lebedev et al. 2005) showed that, in a center-out manual reaching task, the decoding performance of a neuronal ensemble remained high and stable while more than 50% of single neurons significantly changed their firing rates and variability rankings. Rokni and his colleagues (Rokni, Richardson et al. 2007) had monkeys perform a familiar center-out task manually, and found that more than 60% of the cells changed their preferred direction, modulation depth, or offset of the tuning curve between blocks 1 and 3, while the monkeys maintained decent performance. These changes seemed to be random as they did not correlate between cells. Rokni then proposed a redundant network model to simulate the task, and found that local noise and high redundancy could explain how the population of cells maintains performance while individual cells’ tuning curves change randomly.

## Neural Code for BMI

Tehovnik quoted complexities of neuronal encoding of motor commands (e.g., dependence of directional tuning on the arm starting position) as an obstacle to BMI development. In their opinion, BMIs cannot be efficient unless we understand precisely how motor cortical neurons represent movements. We disagree with this statement. The BMI research helps to increase our understanding of brain mechanisms (Nicolelis and Lebedev 2009). Advanced decoding methods, such as the unscented Kalman filter (Li, O’Doherty et al. 2009) and recalibrated feedback intention-trained Kalman filter (Gilja, Nuyujukian et al. 2012) successfully incorporate motor complexities and outperform simpler decoders. Additionally, adaptive filters can compensate for dynamical changes in neuronal tuning properties (Taylor, Tillery et al. 2002, Li, O’Doherty et al. 2011, Sussillo, Stavisky et al. 2016).

## Suppressing Body Movements during BMI

Tehovnik claimed that eye and body movements are integrally linked to neural encoding, and their suppression may decrease the accuracy of BMI control. While this statement merits experimental validation, it has been already shown that subjects can improve in BMI control without producing overt movements with their limbs (Ganguly, Dimitrov et al. 2011). Such improvements have been documented in many studies. For example, in our study of a bimanual BMI (Ifft, Shokur et al. 2013), we eliminated the use of the joystick during training. We had monkeys learn complex bimanual operations from passive observations of the movements performed by two avatar arms. EMG recordings confirmed that the monkeys did not assist themselves with their own arm movements during either the passive observation period or BMI operations. Yet, BMI control performance improved steadily from session to session until reaching levels compatible with the best results obtained so far with BMIs.

## Role of Vision

Similar to the point regarding the interference of body movements with BMI control, Tehovnik expressed a concern regarding the interference of eye movements. While this concern is valid, one should not underestimate the brain’s ability to independently control the eyes, arms and spatial attention (Lebedev and Wise 2001, Lebedev, Messinger et al. 2004).

## Conclusions

Our examination of the claims made by Tehovnik and his colleagues regarding the current state of the BMI field showed several flaws. We support, in principle, the idea of using ITR as a universal metric to compare different BMIs reported in the literature. However, such a comparison should be based on correct math.

Curiously, despite the incorrect assessment of ITR, the main point of Tehovnik and his colleagues is correct that ITR of BMIs is low compared to manual control; even though more recent BMI studies have reported relatively high ITRs (Kao, Nuyujukian et al. 2017, Pandarinath, Nuyujukian et al. 2017). But what is the big surprise? This result is expected for the simple reason that BMIs utilize a miniscule proportion of neurons (hundreds) compared to the total number of neurons involved in normal control (hundreds of thousands or millions, depending on the brain structure used for neural decoding). Yet, Tehovnik and his colleagues missed the key point: despite such a small neuronal sample, monkeys and now human subjects have been able to perform motor tasks of real significance using such an approach. At this juncture, it is reasonable to ask the most basic question: In the future, would patients even care about the ITR if they can benefit from using a neuroprosthetic device built around the BMI concept to aid recovery of basic motor behaviors that could not be generated otherwise? The metric for BMI success in this scenario is certainly not ITR per se, but how much improvement in quality of life that BMIs may provide to a population who have very few really effective therapeutic options available to them. The reality, based upon facts, is that monkeys using BMIs can perform complex motor tasks and the growing number of clinical studies using BMIs are showing that patients can extract significant benefits from them as well. To determine how well a particular BMI type performs, therefore, we should consider the real needs of the patients for whom BMIs are intended (Serruya 2014). Is the primary goal of those with arm paralysis to be able to play a violin or regain some basic motor skills, such as the ability to feed themselves, hold a car key and steer a car? If basic motor skills are primarily needed, and a slower performance speed can be tolerated, then a drop from 5 bits/s to, say, 0.5 bits/s may be not as dramatic or relevant as Tehovnik would like to suggest.

In several places in their article, Tehovnik and his colleagues implied that the “good old days” methodology is superior to the current BMI approach where BMI researchers are attempting to record neurons in large numbers indiscriminately. Would a single electrode precisely placed by an experienced neurophysiologist do a better job to restore mobility in a paraplegic patient? The answer to this question is a no. The mention of excellent task-related neurons recorded by experts in neurophysiology seems credible, but this kind of argument is often based on a statistical manipulation where good recording days are overemphasized, and numerous failures are discarded during every recording session. These details cannot escape scrutiny and recalibration by experienced neurophysiologists who have seen both the past and the present standards of primate cortical neurophysiology. Any experienced neurophysiologist knows about unsuccessful recording sessions when nothing was recorded during the entire day in which some microelectrode could not even penetrate through the animal’s dura. Incidentally, it is untrue that our laboratory does not use movable electrodes. We have developed a 1,000-channel recording system where all microelectrodes are movable (Schwarz, Lebedev et al. 2014). Different from Tehovnik et al. we believe that different recording techniques are appropriate for approaching distinct experimental questions. A single movable microelectrode can work fine in a neurophysiological study where an investigator needs to hold a neuron for 1 hour. But BMI research has different demands, and the technology that we currently have available has evolved with these demands in mind.

Several critical points made by Tehovnik have been already clarified in previous reviews of the BMI field (Lebedev and Nicolelis 2006, Nicolelis and Lebedev 2009). In particular, we have already explained why more neurons are better for BMIs and why the idea of “phenomenal” single cells is impractical. The proposal to use just a few neurons for BMI control is not new, and even the original proponents of this idea have already abandoned this argument and moved toward large-scale neural population recordings.

Another point of Tehovnik is the suggestion that we should investigate brain mechanisms, get a better understanding of how the brain works (Baranauskas 2014) and only then attempt to reproduce brain function with a BMI. This recommendation implies that a better understanding of brain mechanisms can be only gained with the old methods of single electrode recordings, and technological advances made by modern BMIs are of no help. We are puzzled why Tehovnik and his co-authors are so convinced that BMI research prevents endeavors to understand the brain better and that the only way to continue such endeavors is to halt BMI work. In response to this argument, we should ask whether therapeutic techniques such as deep brain stimulation, for which no mechanism has yet been identified, should simply be abandoned until a clear mechanistic explanation is found. What would the more than 100,000 Parkinsonian patients who benefit from this therapy have to say about that?

Among the issues that need to be understood before BMI could become valuable, Tehovnik mentions neuronal directional tuning and brain mechanisms of eye movements. While noting that BMIs could use some of the brain circuitry that normally controls eye movements, Tehovnik does not mention that classical oculomotor studies have a number of problems. In those studies, investigators typically restrained monkey behavior quite drastically (Sparks and Mays 1980, Funahashi, Bruce et al. 1990). Typically, a monkey would be required to stare at the center of the screen for several seconds until it was allowed to change its gaze angle. This highly constrained and overtrained behavior serves the purpose to make an experiment “controlled”, but results in neuronal patterns, which are different from brain activities that enable natural eye movement behavior. BMI research is now leading attempts to investigate motor behaviors in unrestrained animals (Schwarz, Lebedev et al. 2014).

Overall, we hope that our commentary to the paper by Tehovnik and his colleagues will be helpful for the development of better information theory approaches to BMIs, the improvement of BMIs, and the advancement of relevant fields of basic neuroscience.

## References

Ajiboye, A. B., F. R. Willett, D. R. Young, W. D. Memberg, B. A. Murphy, J. P. Miller, B. L. Walter, J. A. Sweet, H. A. Hoyen and M. W. Keith (2017). “Restoration of reaching and grasping movements through brain-controlled muscle stimulation in a person with tetraplegia: a proof-of-concept demonstration.” The Lancet 389(10081): 1821–1830.

Baranauskas, G. (2014). “What limits the performance of current invasive brain machine interfaces?” Frontiers in systems neuroscience 8.

Carmena, J. M., M. A. Lebedev, R. E. Crist, J. E. O’Doherty, D. M. Santucci, D. F. Dimitrov, P. G. Patil, C. S. Henriquez and M. A. Nicolelis (2003). “Learning to control a brain-machine interface for reaching and grasping by primates.” PLoS Biol 1(2): E42.

Carmena, J. M., M. A. Lebedev, C. S. Henriquez and M. A. Nicolelis (2005). “Stable ensemble performance with single-neuron variability during reaching movements in primates.” J Neurosci 25(46): 10712–10716.

Collinger, J. L., B. Wodlinger, J. E. Downey, W. Wang, E. C. Tyler-Kabara, D. J. Weber, A. J. McMorland, M. Velliste, M. L. Boninger and A. B. Schwartz (2013). “High-performance neuroprosthetic control by an individual with tetraplegia.” Lancet 381(9866): 557–564.

Fitts, P. M. and R. L. Deininger (1954). “S-R compatibility: correspondence among paired elements within stimulus and response codes.” J Exp Psychol 48(6): 483–492.

Fitts, P. M. and J. R. Peterson (1964). “Information Capacity of Discrete Motor Responses.” J Exp Psychol 67: 103–112.

Fitzsimmons, N. A., M. A. Lebedev, I. D. Peikon and M. A. Nicolelis (2009). “Extracting kinematic parameters for monkey bipedal walking from cortical neuronal ensemble activity.” Front Integr Neurosci 3: 3.

Funahashi, S., C. J. Bruce and P. S. Goldman-Rakic (1990). “Visuospatial coding in primate prefrontal neurons revealed by oculomotor paradigms.” Journal of neurophysiology 63(4): 814–831.

Ganguly, K. and J. M. Carmena (2009). “Emergence of a stable cortical map for neuroprosthetic control.” PLoS Biol 7(7): e1000153.

Ganguly, K., D. F. Dimitrov, J. D. Wallis and J. M. Carmena (2011). “Reversible large-scale modification of cortical networks during neuroprosthetic control.” Nature neuroscience 14(5): 662–667.

Georgopoulos, A. P. and J. T. Massey (1988). “Cognitive spatial-motor processes. 2. Information transmitted by the direction of two-dimensional arm movements and by neuronal populations in primate motor cortex and area 5.” Exp Brain Res 69(2): 315–326.

Gilja, V., P. Nuyujukian, C. A. Chestek, J. P. Cunningham, B. M. Yu, J. M. Fan, M. M. Churchland, M. T. Kaufman, J. C. Kao, S. I. Ryu and K. V. Shenoy (2012). “A high-performance neural prosthesis enabled by control algorithm design.” Nat Neurosci 15(12): 1752–1757.

Hochberg, L. R., D. Bacher, B. Jarosiewicz, N. Y. Masse, J. D. Simeral, J. Vogel, S. Haddadin, J. Liu, S. S. Cash and P. van der Smagt (2012). “Reach and grasp by people with tetraplegia using a neurally controlled robotic arm.” Nature 485(7398): 372.

Huber, D., D. A. Gutnisky, S. Peron, D. H. O’Connor, J. S. Wiegert, L. Tian, T. G. Oertner, L. L. Looger and K. Svoboda (2012). “Multiple dynamic representations in the motor cortex during sensorimotor learning.” Nature 484(7395): 473–478.

Ifft, P. J., S. Shokur, Z. Li, M. A. Lebedev and M. A. Nicolelis (2013). “A brain-machine interface enables bimanual arm movements in monkeys.” Sci Transl Med 5(210): 210ra154.

Kao, J. C., P. Nuyujukian, S. I. Ryu and K. V. Shenoy (2017). “A high-performance neural prosthesis incorporating discrete state selection with hidden Markov models.” IEEE Transactions on Biomedical Engineering 64(4): 935–945.

Lebedev, M. A. (2014). “How to read neuron-dropping curves?” Front Syst Neurosci 8: 102.

Lebedev, M. A., A. Messinger, J. D. Kralik and S. P. Wise (2004). “Representation of attended versus remembered locations in prefrontal cortex.” PLoS Biol 2(11): e365.

Lebedev, M. A. and M. A. Nicolelis (2006). “Brain-machine interfaces: past, present and future.” Trends Neurosci 29(9): 536–546.

Lebedev, M. A. and M. A. Nicolelis (2011). “Toward a whole-body neuroprosthetic.” Prog Brain Res 194: 47–60.

Lebedev, M. A. and M. A. Nicolelis (2017). “Brain-Machine Interfaces: From Basic Science to Neuroprostheses and Neurorehabilitation.” Physiological Reviews 97(2): 767–837.

Lebedev, M. A. and S. P. Wise (2001). “Tuning for the orientation of spatial attention in dorsal premotor cortex.” Eur T Neurosci 13(5): 1002–1008.

Li, Z., J. E. O’Doherty, T. L. Hanson, M. A. Lebedev, C. S. Henriquez and M. A. Nicolelis (2009). “Unscented Kalman filter for brain-machine interfaces.” PLoS One 4(7): e6243.

Li, Z., J. E. O’Doherty, M. A. Lebedev and M. A. Nicolelis (2011). “Adaptive decoding for brain-machine interfaces through Bayesian parameter updates.” Neural Comput 23(12): 3162–3204.

Mulliken, G. H., S. Musallam and R. A. Andersen (2008). “Decoding trajectories from posterior parietal cortex ensembles.” T Neurosci 28(48): 12913–12926.

Newton, I. (1687). Sir Isaac Newton’s mathematical principles of natural philosophy and his system of the world. Berkeley, University of California.

Nicolelis, M. A. and M. A. Lebedev (2009). “Principles of neural ensemble physiology underlying the operation of brain-machine interfaces.” Nat Rev Neurosci 10(7): 530–540.

O’Doherty, J. E., M. A. Lebedev, P. J. Ifft, K. Z. Zhuang, S. Shokur, H. Bleuler and M. A. Nicolelis (2011). “Active tactile exploration using a brain-machine-brain interface.” Nature 479(7372): 228–231.

Pandarinath, C., P. Nuyujukian, C. H. Blabe, B. L. Sorice, J. Saab, F. R. Willett, L. R. Hochberg, K. V. Shenoy and J. M. Henderson (2017). “High performance communication by people with paralysis using an intracortical brain-computer interface.” eLife 6: e18554.

Pathirana, P., A. Savkin and S. Jha (2004). “Location estimation and trajectory prediction for cellular networks with mobile base stations.” IEEE Trans Veh Technol 53: 1903–1913.

Pierce, J. R. (1980). An introduction to information theory: symbols, signals & noise. New York, Dover Publications.

Rokni, U., A. G. Richardson, E. Bizzi and H. S. Seung (2007). “Motor learning with unstable neural representations.” Neuron 54(4): 653–666.

Rosales, R. and S. Sclaroff (1999). “Trajectory guided recognition of actions.” P Soc Photo-Opt Ins 3840: 25–36.

Sakitt, B. (1980). “Visual-Motor Efficiency (VME) and the Information Transmitted in Visual-Motor Tasks.” B Psychonomic Soc 16: 329–332.

Schmidt, E. M., J. S. McIntosh, L. Durelli and M. J. Bak (1978). “Fine control of operantly conditioned firing patterns of cortical neurons.” Exp Neurol 61(2): 349–369.

Schwarz, D. A., M. A. Lebedev, T. L. Hanson, D. F. Dimitrov, G. Lehew, J. Meloy, S. Rajangam, V. Subramanian, P. J. Ifft, Z. Li, A. Ramakrishnan, A. Tate, K. Z. Zhuang and M. A. Nicolelis (2014). “Chronic, wireless recordings of large-scale brain activity in freely moving rhesus monkeys.” Nat Methods 11(6): 670–676.

Serruya, M. (2014). “Bottlenecks to clinical translation of direct brain-computer interfaces.” Front Syst Neurosci.

Shannon, C. and W. Weaver (1964). The mathematical theory of communication. Urbana, Illinois, University of Illinois Press.

Sparks, D. L. and L. E. Mays (1980). “Movement fields of saccade-related burst neurons in the monkey superior colliculus.” Brain research 190(1): 39–50.

Sussillo, D., S. D. Stavisky, J. C. Kao, S. I. Ryu and K. V. Shenoy (2016). “Making brain-machine interfaces robust to future neural variability.” Nature communications 7.

Taylor, D. M., S. I. H. Tillery and A. B. Schwartz (2002). “Direct cortical control of 3D neuroprosthetic devices.” Science 296(5574): 1829–1832.

Tehovnik, E. J., L. C. Woods and W. M. Slocum (2013). “Transfer of information by BMI.” Neuroscience 255: 134–146.

Wessberg, J., C. R. Stambaugh, J. D. Kralik, P. D. Beck, M. Laubach, J. K. Chapin, J. Kim, S. J. Biggs, M. A. Srinivasan and M. A. Nicolelis (2000). “Real-time prediction of hand trajectory by ensembles of cortical neurons in primates.” Nature 408(6810): 361–365.

Wolpaw, J. R., N. Birbaumer, W. J. Heetderks, D. J. McFarland, P. H. Peckham, G. Schalk, E. Donchin, L. A. Quatrano, C. J. Robinson and T. M. Vaughan (2000). “Brain-computer interface technology: a review of the first international meeting.” IEEE Trans Rehabil Eng 8(2): 164–173.

Wolpaw, J. R., N. Birbaumer, D. J. McFarland, G. Pfurtscheller and T. M. Vaughan (2002). “Brain-computer interfaces for communication and control.” Clin Neurophysiol 113(6): 767–791.

Wolpaw, J. R. and D. J. McFarland (2004). “Control of a two-dimensional movement signal by a noninvasive brain-computer interface in humans.” Proc Natl Acad Sci U S A 101(51): 17849–17854.

